# Optimizing Cryo-EM Structural Analysis of G_i_-coupling Receptors via Engineered G_t_ and Nb35 Application

**DOI:** 10.1101/2023.10.07.561347

**Authors:** Hidetaka S. Oshima, Fumiya K. Sano, Hiroaki Akasaka, Aika Iwama, Wataru Shihoya, Osamu Nureki

## Abstract

Cryo-EM single particle analysis has recently facilitated the high-resolution structural determination of numerous GPCR-G complexes. Diverse methodologies have been devised with this trend, and in the case of GPCR-G_i_ complexes, scFv16, an antibody that recognizes the intricate interface of the complex, has been mainly implemented to stabilize the complex. However, owing to their flexibility and heterogeneity, structural determinations of GPCR-G_i_ complexes remain both challenging and resource-intensive. By employing eGα_t_, which exhibits binding affinity to modified nanobody Nb35, the cryo-EM structure of Rhodopsin-eGα_t_ complex was previously reported. Using this modified G protein, we determined the structure of the ET_B_-eG_t_ complex bound to the modified Nb35. The determined structure of ET_B_ receptor was the same as the previously reported ET_B_-G_i_ complex, and the resulting dataset demonstrated significantly improved anisotropy. This modified G protein will be utilized for the structural determination of other GPCR-G_i_ complexes.

**Highlights:**

- The study introduces the engineered G protein subunit eGα_T_, which enhances the resolution of GPCR-G protein structures by suppressing G protein conformational fluctuations and is particularly beneficial for G_i_-coupled receptors.
- The cryo-EM structure of the ET_B_ receptor complexed with eG_t_-Nb35 reveals improved map quality, reduced anisotropy, and isotropic density distribution, increasing the accuracy of structural analysis.
- Structural comparison between ET_B_-G_i_ and ET_B_-eG_t_ reveals similar receptor-G protein interactions, demonstrating the utility of eG_t_-Nb35 for studying GPCR-G_i_ complexes and the potential for broader applications within the G_i_ family.

## Introduction

G protein-coupled receptors (GPCRs) represent the most expansive superfamily within the human system. They have engendered profound research scrutiny, primarily attributable to their critical roles in cellular communication and their potential as therapeutic targets. Numerous cryo-electron microscopy (cryo-EM) structures of GPCR-G protein complexes have recently been reported^1,2^, effectively catalyzing a paradigm shift in drug design strategies^3^. One of the factors behind this success is the development of structure-recognizing antibodies for G proteins. Nanobody 35 (Nb35) is an antibody that specifically recognizes the interface between the RAS-like domain of Gα_s_ and Gβ^4^. Notably, Nb35 inhibits the GTP-GDP exchange reaction, consequently stabilizing the GPCR-G protein complex in the most stable nucleotide-free conformation, and thus allowing high resolution structure determination^2,5–9^. The single-chain variable fragment of mAb16 (scFv16), another remarkable antibody, specifically recognizes the interface between the N-terminal helix of Gα_i_ (αN helix) and Gβ^10^. scFv16 also imparts resistance to GTPγS on the G protein and enhances the requisite features for cryo-EM structural analysis, thereby facilitating structural determination^1,2,11–14^. Furthermore, the versatile applicability of scFv16 extends to other G-proteins through substitution with the αN helix of the Gα subunit.

However, challenges remain in the structural analyses of GPCR-G complexes, especially G_i_. scFv16 weakly inhibits the GDP/GTP exchange reaction, in stark contrast to the robust inhibition conferred by Nb35^4,10^. Furthermore, scFv16 binds to the N-terminal helix of G_i_, as opposed to the RAS-like domains of Gαs and Gβ, thereby exerting a somewhat attenuated conformational stabilizing effect upon the G protein. These features lead to structural polymorphism of the receptor-G-protein interface, as observed in the LPA1-G_i_ complex^12^. Additionally, the binding of scFv16 imparts a flattening effect on the GPCR-G protein complex, which results in orientation bias, as unequivocally evident in the representation of 2D class averages for typical GPCR-G_i_ complexes. For μOR-G_i_, improved resolution by masking scFv16 during data processing has been reported, indicating that scFv16 binding is not rigid^15^. Even though scFv16 facilitates the complex formation and particle alignment during the cryo-EM data processing, the G_i_ complex structural analysis requires a relatively large data set, compared to the G_s_-coupled receptors.

One of the solutions to this problem is the use of eGα_T_^16^. The native Nb35 cannot bind to the visual Rho-G_t_ complex formed with rGα_T_. In contrast, the engineered Nb35, termed Nb35∗, binds to the Rho-G_t_ complex formed with the engineered rGα_T_ subunit, termed eGα_T_. Cryo-EM structures of Rho-eGα_T_, in both the presence and absence of Nb35*, revealed that Nb35* improved the resolution of the complex structure by suppressing the G protein conformational fluctuations. These studies inspired us to apply eGα_T_-Nb35* to the structural analysis of other G_i_-coupled receptors, because G_t_ belongs to the G_i_ family. Here, we report the cryo-EM structure of the ET_B_-receptor-eG_t_-Nb35* complex, thereby underscoring the potential utility of eGα_T_-Nb35*.

## Materials and Methods

### Constructs

The sequence of eG_t_ was based on the previously reported eGα_T_^16^, but the αN domain was replaced with that of human G_i1_. eG_t_ was fused to the C-terminus of bovine Gγ_2_, following the GSAGSAGSA linker. Rat Gβ_1_ was cloned with a C-terminal HiBiT connected with the 15 amino sequence GGSGGGGSGGSSSGG, as described previously^13^. The resulting Gγ_2_-eG_t_ and Gβ_1_-HiBiT constructs were subcloned into the pFastBac Dual vector, as described previously^7^. The full-length human ET_B_ gene was subcloned into the pFastBac vector, with the native signal peptide replaced with the HA-signal peptide and the DYKDDDDK Flag epitope tag. LgBiT was fused to the C-terminus of the ET_B_ gene, followed by a 3C protease site and an eGFP-His8 tag. GGSGGGGSGGSSSGG linkers were inserted on both the N-terminal and C-terminal sides of LgBiT.

### ET-1–ET_B_– eG_t_ complex preparation

For expression, 1 L of Sf9 cells at a density of 3 × 10^6^ cells/ml were co-infected with baculovirus encoding the ET_B_-LgBiT-eGFP and eG_t_ trimer at the ratio of 1:1. Cells were harvested 48 h after infection. Cell pellets were resuspended and Dounce-homogenized in 20 mM Tris-HCl, pH 8.0, 100 mM NaCl, and 10% glycerol. Apyrase was added to the lysate at a final concentration of 25 mU/ml, and ET-1 was added at a final concentration of 2 µM. The lysate was incubated at room temperature for 1 h. Then, the membrane fraction was collected by ultracentrifugation at 180,000*g* for 1 h. The cell membrane was solubilized in buffer, containing 20 mM Tris-HCl, pH 8.0, 150 mM NaCl, 1% DDM, 0.2% CHS, 10% glycerol, and 2 μM ET-1, for 1 h at 4 °C. The supernatant was separated from the insoluble material by ultracentrifugation at 180,000*g* for 30 min and then incubated with the Anti-DYKDDDDK M2 resin (Genscript) for 1 h. The resin was washed with 20 column volumes of wash buffer, containing 20 mM Tris-HCl, pH 8.0, 500 mM NaCl, 10% glycerol, 0.1% Lauryl Maltose Neopentyl Glycol (LMNG; Anatrace), and 0.01 % CHS. The complex was eluted with buffer, containing 20 mM Tris-HCl, pH 8.0, 150 mM NaCl, 10% glycerol, 0.1% LMNG, 0.01% CHS, and 0.15 mg/ml Flag peptide. The eluate was treated with 0.5 mg of HRV-3C protease (prepared in-house) and 0.5 mg of Nb35*, prepared as described previously^16^. The complex was concentrated and separated by size exclusion chromatography on a Superdex 200 Increase 10/300 column, in 20 mM Tris-HCl, pH 8.0, 150 mM NaCl, 0.01% LMNG, 0.001% CHS, and 1 μM agonist. Peak fractions were concentrated for the cryo-EM grid preparation.

### Cryo-EM grid preparation and data acquisition

Three μL of the purified complex were applied onto a freshly glow-discharged Quantifoil holey carbon grid (R1.2/1.3, Au, 300 mesh), which was then plunge-frozen in liquid ethane by using a Vitrobot Mark IV (FEI). Cryo-EM data collection was performed on a 300 kV Titan Krios G3i microscope (Thermo Fisher Scientific) equipped with a BioQuantum K3 imaging filter (Gatan) and a K3 direct electron detector (Gatan). A total of 5,253 movies were acquired with a calibrated pixel size of 0.83 Å pix−1 and a defocus range of −0.8 to −1.6 μm, using the EPU software (Thermo Fisher Scientific’s single-particle data collection software). Each movie was acquired for 2.3 s and split into 48 frames, resulting in an accumulated exposure of about 48 e^−^ Å^−2^.

All acquired movies in the super-resolution mode were binned by 2× and then dose-fractionated and subjected to beam-induced motion correction implemented in RELION 3.1^17^. The contrast transfer function (CTF) parameters were estimated using patch CTF estimation in cryoSPARC v3.3^18^. Particles were initially picked from a small fraction of the micrographs with the Blob picker, and subjected to several rounds of two-dimensional (2D) classification in cryoSPARC. Selected particles were used for training the Topaz model^19^. For the full dataset, 3,359,173 particles were picked and extracted with a pixel size of 3.32 Å. The particles went through several rounds of hetero-refinement to remove trash classes. The resulting particles were re-extracted with a pixel size of 1.29 Å and curated by three-dimensional (3D) classification without alignment in RELION. Consequently, the 52,285 particles in the best class were reconstructed using non-uniform refinement, resulting in a 3.24 Å overall resolution reconstruction, with the gold standard Fourier Shell Correlation (FSC=0.143) criteria in cryoSPARC. After that, the 3D model was refined with masks on the receptor and the G protein trimer. The local refined maps were composited using phenix.combine_focused_maps and used for modeling^20^. The workflow of the cryo-EM analysis is summarized in Supplementary Fig. 1b.

**Figure 1.**
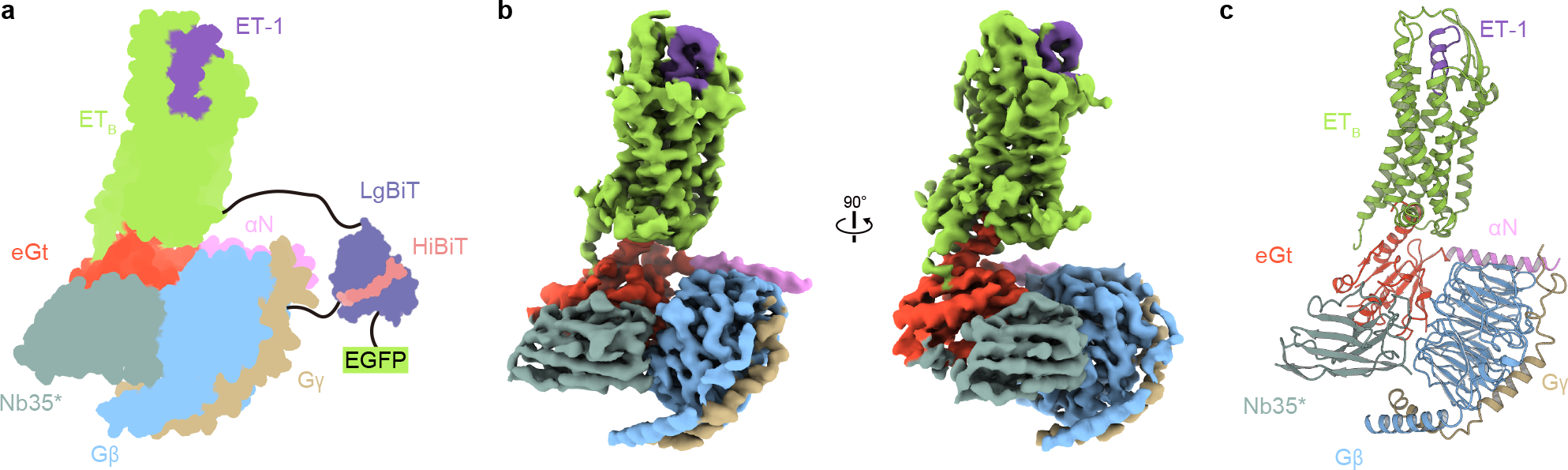
Overall structure of the ET-1-ET_B_-eG_t_ complex. (a) Schematic representation of the ET_B_-eG_t_ complex. (b) Cryo-electron microscopy (cryo-EM) map, with each component colored differently. (c) Structure of the complex determined after refinement in the cryo-EM map, shown as a ribbon representation.

### Model building and refinement

The quality of the density map was sufficient to build an atomic model. The previously reported high-resolution crystal structure of the ET-3 bound ET_B_ receptor (PDB 6IGK) and the cryo-EM Rho-G_t_ structure (PDB 6OYA) were used as the initial models for the model building of the receptor and eG_t_ portions, respectively^16,21^. At first, the models were fitted into the density map by jiggle fit using COOT, and then the atomic models were readjusted into the density map using COOT and refined using phenix.real_space_refine (v1.19) with the secondary structure restraints, using phenix.secondary_structure_restraints^20,22,23^.

## Results

### Structure determination

For the structural analysis employing eG_t_-Nb35*, we chose the human endothelin ET_B_ receptor that recognizes the vasoactive peptide endothelin-1 (ET-1). Significantly, the structural information on ET_B_ is quite comprehensive among GPCRs^13,21,24–28^. In 2023, we reported the cryo-EM structure of the ET_B_-G_i_ complex by the Fusion-G system^13^, which seamlessly amalgamates two complex stabilization methodologies; namely, the nanobit-tethering strategy and G_i_-Gγ fusion. However, the presence of orientation bias necessitated the acquisition of an extensive dataset of approximately 10,000 movies. Remarkably, the persistence of orientation bias remained evident in the final map. By comparing this study of the ET_B_-G_i_ complex, we attempted to evaluate the structural analysis with eG_t_-Nb35*.

The design of eG_t_ was based on eGα_T_^16^, whose αN domain was replaced with that of human G_i1_ (Fig. 1a). This modification allowed for the additional employment of scFv16, although we did not use it for this experiment. The gene encoding eG_t_ was cloned into a 3-in-1 vector for G-protein trimer expression, in which the Gα subunit was fused to the C-terminus of the Gγ subunit. The ET_B_-LgBiT fusion was used as described previously^13^. ET_B_-LgBiT and the G-protein trimer were co-expressed in insect cells, and subsequently the ET_B_-eG_t_ complex was purified by Flag affinity chromatography. After an incubation with Nb35*, the complex was separated by size exclusion chromatography and concentrated for grid preparation (Supplementary Fig. 1a). Finally, we determined the cryo-EM structure of the ET_B_-eG_t_ complex with an overall resolution of 3.2 Å (Fig. 1b, 1c, Supplementary Fig. 1b). Notably, the representative classes derived from 2D classification exhibited a broader range of orientations compared to those observed for the ET_B_-G_i_ complex (Fig. 2a and 2b). We further refined the density map using masks focused on the receptor and the G protein. The composite map was used for modeling. Every component, including ET-1, was well resolved.

**Figure 2.**
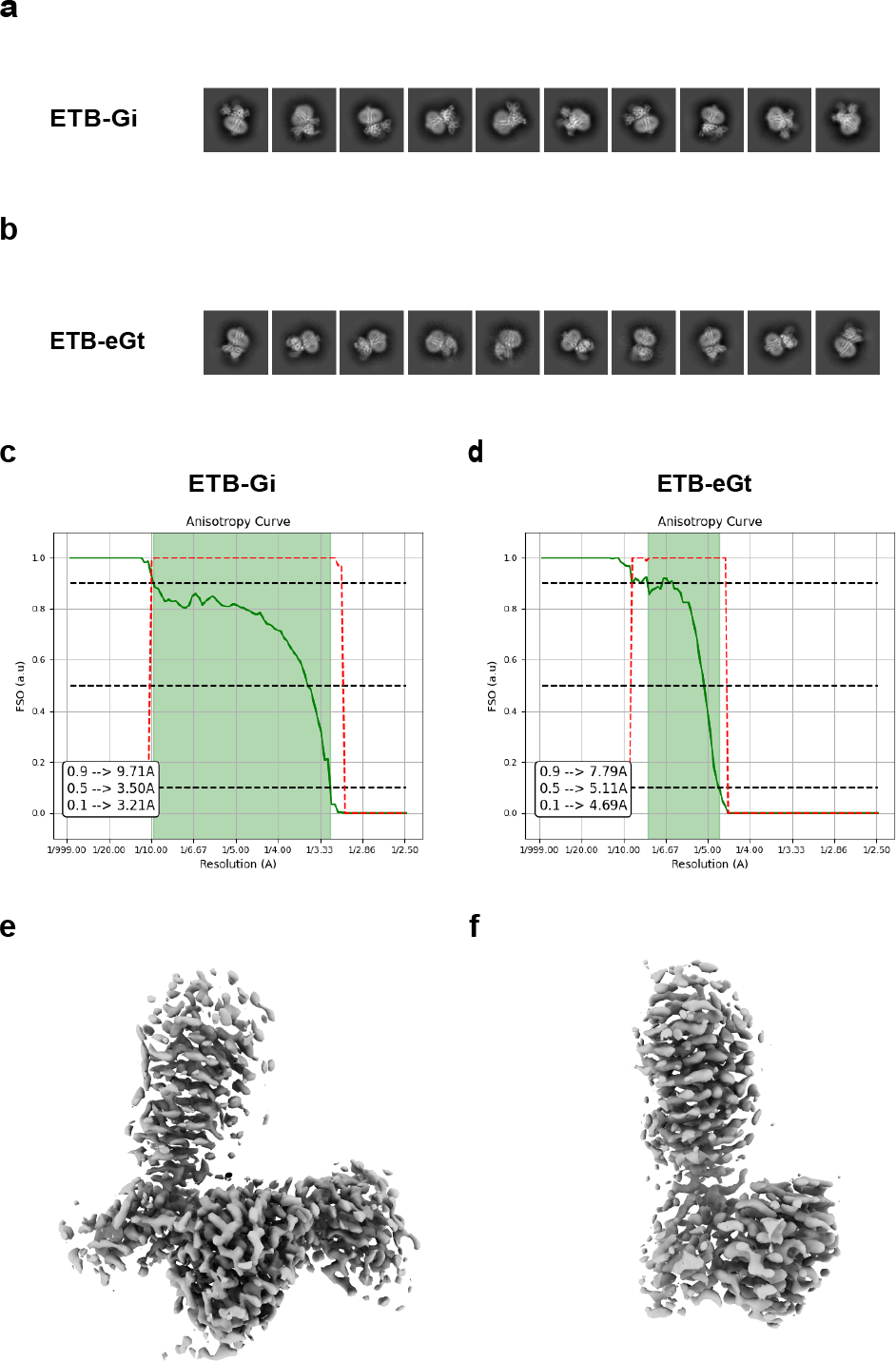
Anisotropy analysis of the ET_B_-G_i_ complex and ET_B_-eG_t_ complex. (a, b) Representative 2D classes of the ETB-G_i_ complex (a) and ETB-eG_t_ complex (b). (c, d) FSO curves of the ET_B_-G_i_ complex (c) and ET_B_-eG_t_ complex (d). (e, f) Subtracted map of the original map and the low pass filtered map of the ET_B_-G_i_ complex (e) and ET_B_-eG_t_ complex (f).

### Improvement of map bias

We evaluated the obtained map by Fourier Shell Correlation (FSC) and assessed its quality with the Fourier Shell Occupancy (FSO) metric^29^. FSO quantifies the proportion of directional FSCs that exceed a threshold of 0.143. This metric provides a concise summary of the reconstruction quality, pinpointing regions where the reconstruction is isotropic and the areas transitioning to anisotropy, and presenting the global resolution in a single plot, enhancing its interpretability.

We determined the FSO values for the overall maps of the ET_B_-G_i_ complex and ET_B_-eG_t_ complex (Fig. 2c and 2d). For ET_B_-G_i_, the FSO value gradually decreased to 0.5 before experiencing a sharp decline (Fig. 2c). In contrast, the FSO value for eG_t_ dropped precipitously (Fig. 2d). Based on these FSO values, we inferred a significant enhancement in the isotropy of the map.

To provide a clearer visual representation of this improvement, we generated subtracted maps (Fig. 2e and 2f). These maps were produced by subtracting the original map from the map that was low pass filtered at the resolution corresponding to an FSO value of 0.5. A map with pronounced anisotropy will result in a subtracted map that appears elongated and continuous^29^. The subtracted map for G_i_ revealed an elongated continuous density, particularly around the G protein region (Fig. 2e). Conversely, the subtracted map for eG_t_ exhibited a more fragmented density pattern (Fig. 2f). These visual cues corroborated our assertion that the map’s anisotropy has been ameliorated.

### Structural comparison with ET_B_-G_i_ and ET_B_-eG_t_

The structures of the ET_B_-G_i_ and ET_B_-eG_t_ receptors superimposed well, with a root mean square deviation of 0.93Å (Fig. 3a). Moreover, the relative G-protein positions were also similar. Subsequently, we focused on the details of the receptor-G-protein interactions at α5H and its underlying hydrophobic pocket, which are essential interaction sites^30^ (Fig. 3c–f). The amino acid residues of α5H do not differ significantly between eG_t_ and G_i_, except for two residues (Fig. 3b). Consistently, the hydrogen-bonding interactions are conserved, especially those encompassing R^3.50^ (superscripts indicate Ballesteros–Weinstein numbers^30^) and the carboxyl oxygen of C351, among the hallmark features of the GPCR-G_i_ complex^31–33^ (Fig. 3c and 3d). Analogously, W206^ICL2^ is ensconced within a hydrophobic pocket at the root of α5H (Fig. 3e and 3f). Overall, the receptor-G interactions with eG_t_ are nearly equivalent to those of G_i_.

**Figure 3.**
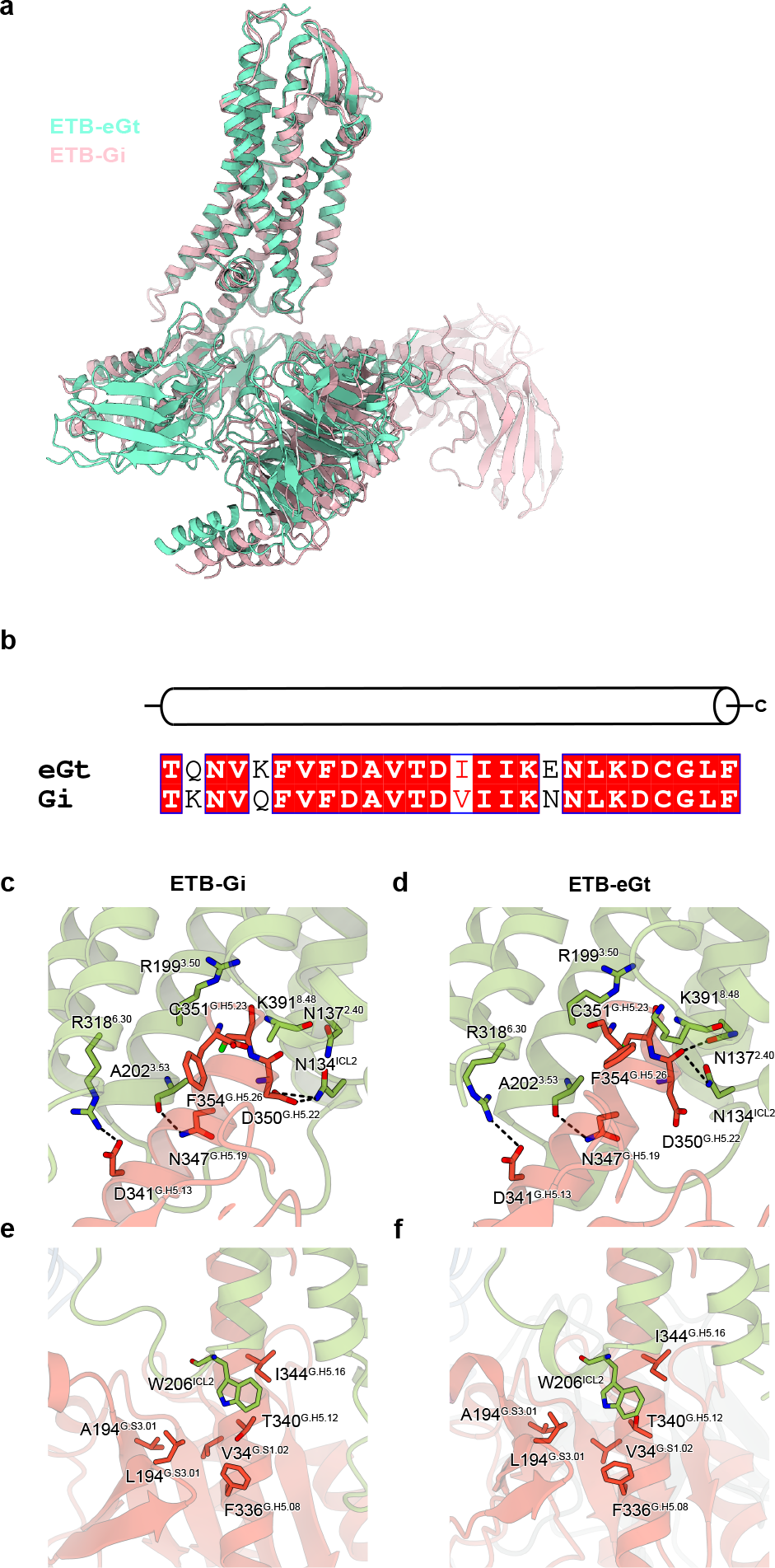
Structural comparison between the ET_B_-G_i_ complex and ET_B_-eG_t_ complex. (a) Superimposition of the ET_B_-G_i_ and ET_B_-eG_t_ complexes, focused on the receptor. (b) Alignment of the amino acid sequences of human G_i1_ and eG_t_, focused on α5H^34^. (c–f) Structural comparison between the ETB-G_i_ (c, e) and ETB-eG_t_ (d, f) complexes. Detailed views of interactions at α5H (c, d) and ICL2 (e, f).

## Conclusion

Cryo-EM structural analysis of GPCRs has been enhanced by the innovative use of antibodies such as Nb35 and scFv16, significantly accelerating our understanding of the complexity of intracellular signal transduction and boosting drug design efforts. In particular, exploratory studies highlighting the potential applications and nuances of using Nb35* and eGα_T_ in scrutinizing the cryo-EM structure of the ET_B_-eG_t_-Nb35* complex are pushing the limits of optimization strategies for GPCR-G protein complex structure analysis. The finding that the ET_B_-eG_t_ complex can be studied with markedly reduced map anisotropy compared to the ET_B_-G_i_ complex not only demonstrates the effectiveness of our approach, but also sparks hope for refining the structural analyses of other GPCR-G protein complexes.

The similarities in receptor-protein interactions observed between eG_t_ and G_i_, ranging from receptor superposition to the retention of hydrogen bond interactions, highlight the conserved mechanisms and structural parallelism, paving the way for further structural and functional studies of GPCRs. However, it should be noted that although the application of eGα_T_ and Nb35* has improved some structural analysis challenges, notably orientation bias and resolution, the search for a rational and universally applicable methodology in the structure determination of GPCR-G protein complexes is still ongoing.

## Abbreviations

ET_B_: Endothelin receptor type B
NB35: Nanobody35
FSO: Fourier shell occupancy

## Author contributions

H. S. O. purified the receptor and performed the complex formation and grid-preparation, assisted by F. K. S., H. A., W. S. and A. I. With assistance from F. K. S. and H. A., H. S. O. performed the cryo-EM observations, data collection, single-particle analysis, model building, and refinement. H. S. O. and W.S. wrote the manuscript with input from all authors. W.S. and O.N. supervised the research.

## Data Availability

Atomic coordinates for the ET_B_-eG_t_ complex have been deposited in the Protein Data Bank, under accession code XXX. The associated electron microscopy data have been deposited in the Electron Microscopy Database and EMD-XXXX, respectively. All other data are available from the corresponding author upon reasonable request.

**Table 1.**
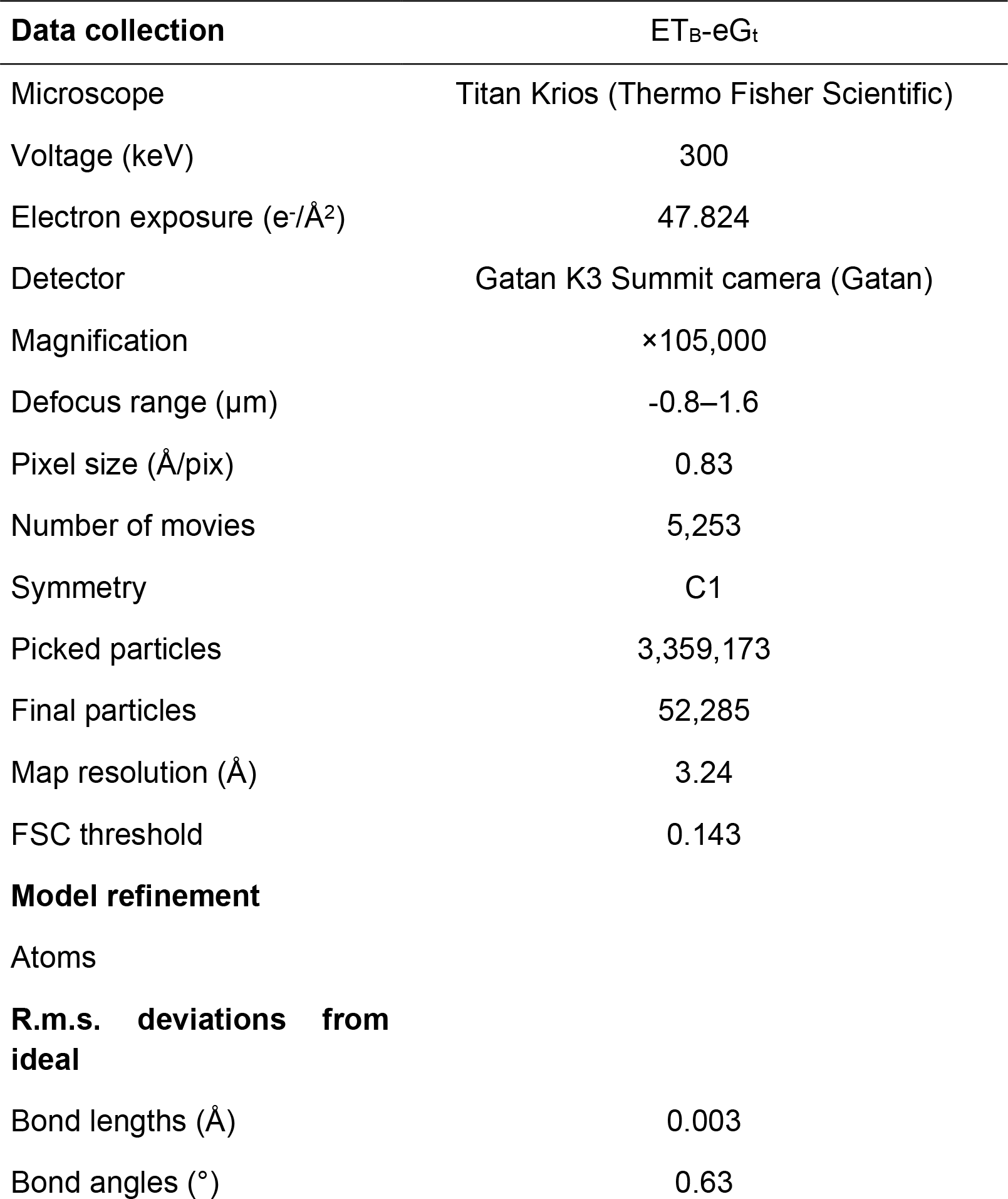
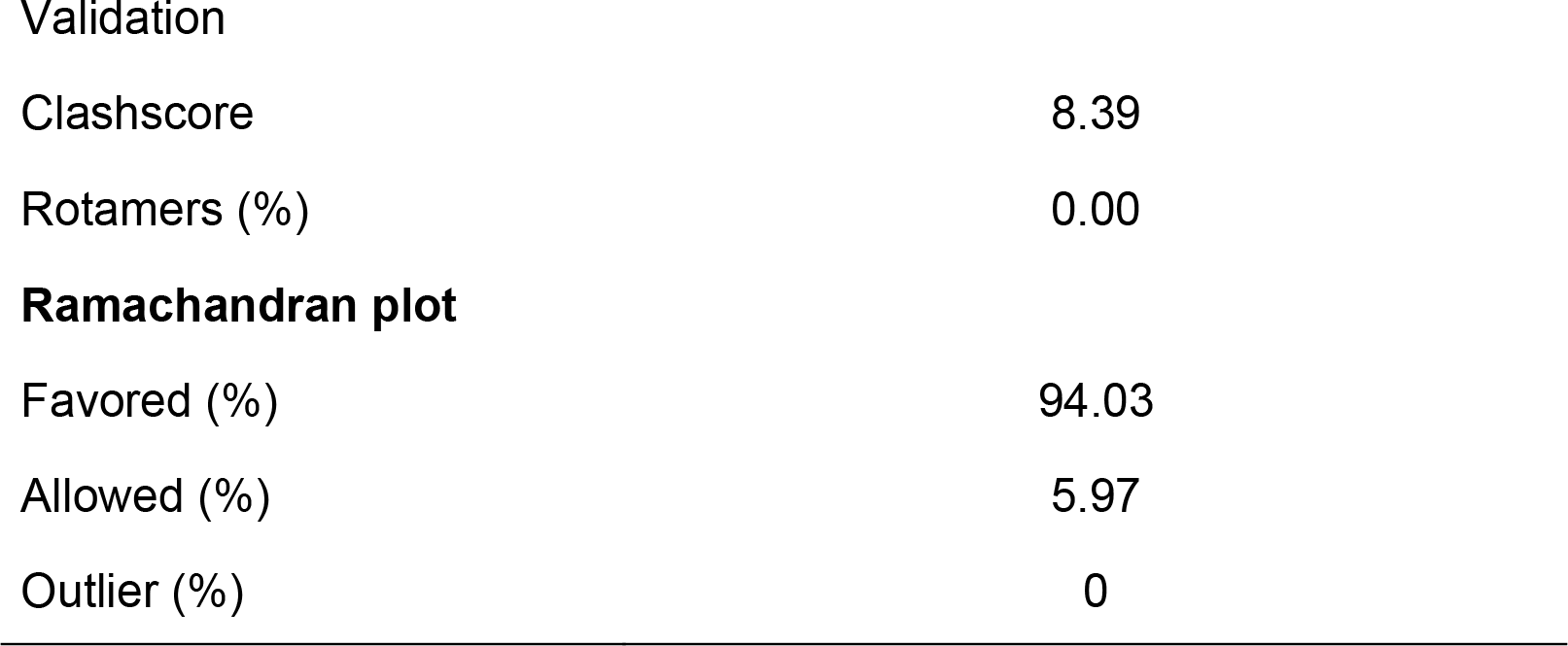

