## Supplementary figures and images for "Optimizing Cryo-EM Structural Analysis of G_i_-coupling Receptors via Engineered G_t_ and Nb35 Application"

### Supplemental Figure 1

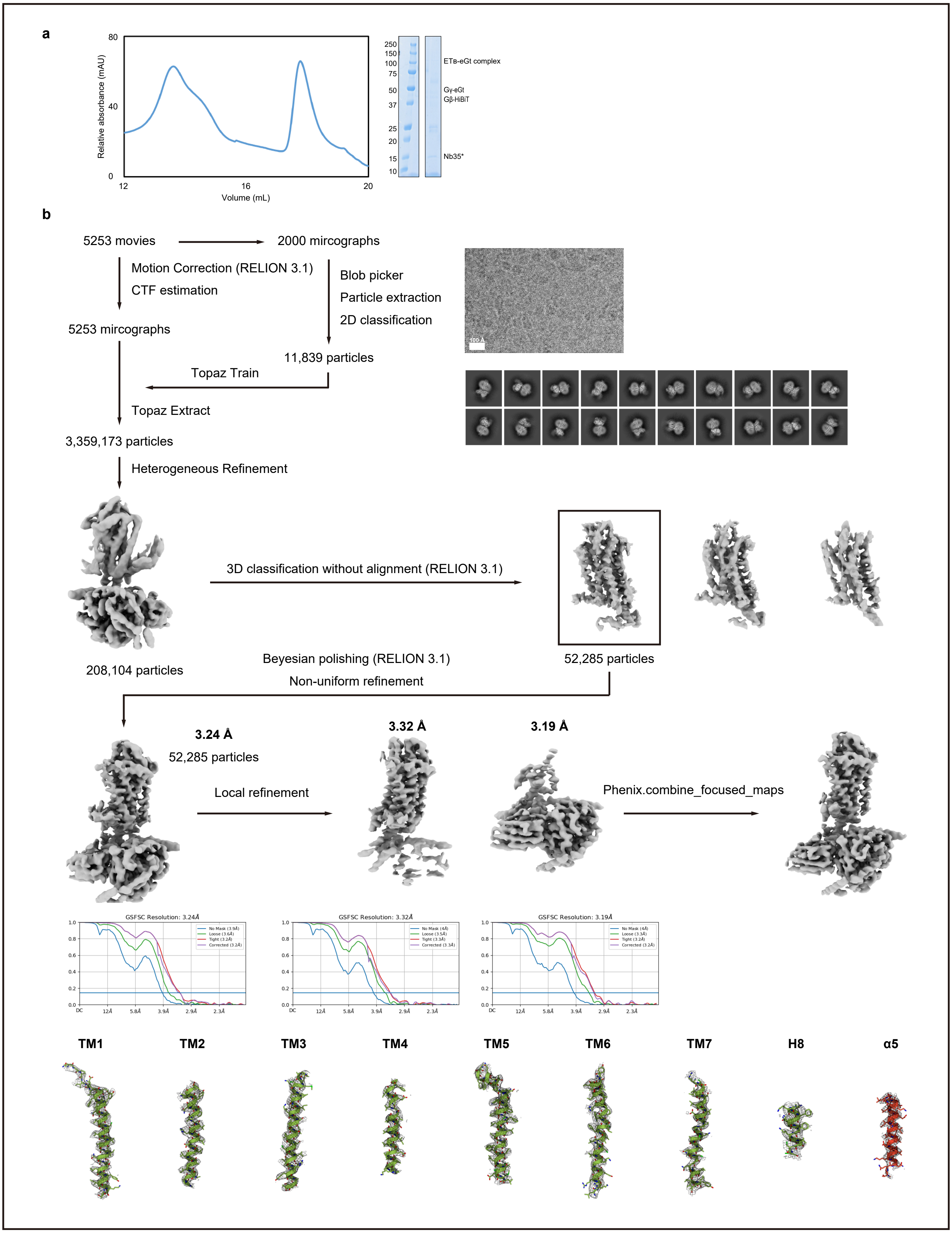


#### Oshima et al. Supplementalfigure 1
